# Establishment of a marine nematode model for animal functional genomics, environmental adaptation and developmental evolution

**DOI:** 10.1101/2020.03.06.980219

**Authors:** Yusu Xie, Pengchi Zhang, Beining Xue, Xuwen Cao, Xiaoliang Ren, Lili Wang, Yuanyuan Sun, Hanwen Yang, Liusuo Zhang

## Abstract

Nematodes play key roles in marine ecosystem. Although oceans cover 71% of the Earth’s surface, none of marine model nematode has been reported. Here, we constructed the first inbred line of free-living marine nematode *Litoditis marina*, sequenced and assembled its genome. Furthermore, we successfully applied CRISPR/Cas9 genome editing in *L. marina*. Comparative genomics revealed that immunity and oxygen regulation genes are expanded, which is probably central to its sediment adaptation. While *L. marina* exhibits massive gene contractions in NHRs, chemoreceptors, xenobiotics detoxification and core histones, which could explain the more defined marine environment. Our experiments showed that dozens of H4 genes in *Caenorhabditis elegans* might contribute to its adaptation to the complex terrestrial environments, while two H4 genes in *L. marina* are involved in salinity stress adaptation. Additionally, ninety-two conserved genes appear to be positively selected in *L. marina*, which may underpin its osmotic, neuronal and epigenetic changes in the sea. With short generation time, highly inbred lines, and genomic resources, our report brings *L. marina* a promising marine animal model, and a unique satellite marine model to the well-known biomedical model nematode *C. elegans*. This study will underpin ongoing work on animal functional genomics, environmental adaptation and developmental evolution.

## INTRODUCTION

Nematodes represent about 80% of all the multicellular animals on Earth, and play major roles in many ecosystems (Eisenhauer and Guerra 2019; van den Hoogen et al. 2019). They occupy essentially all ecological niches: hot springs, polar ice, soil, fresh and salt water, and as parasites of plants, animals and humans. It is reported that 26,646 species have been identified (Hugot et al. 2001; Kikuchi et al. 2017), and the total number of species is estimated to be more than one million (Hugot et al. 2001; Lambshead and Boucher 2003). It is thought that nematodes may have emerged from a marine habitat during the Cambrian Explosion (van den Elsen et al. 2009), and colonized land about 442 million years ago (Rota-Stabelli et al. 2013). There are 11,400 marine nematodes have been described within the phylum of Nematoda (Appeltans et al. 2012), representing approximately 43% of the known nematode species.

As a biomedical model organism, the free-living soil nematode *Caenorhabditis elegans* is the first model nematode, and the first eukaryotic animal ever had its whole genome sequenced (The Caenorhabditis elegans Sequencing Consortium 1998). Other free-living nematodes had their genome sequenced subsequently in the last two decades (Stein et al. 2003; Dieterich et al. 2008; Kanzaki et al. 2018; Ren et al. 2018; Yin et al. 2018; Stevens et al. 2019; Weinstein et al. 2019). Aside from *C. elegans, Pristionchus pacificus* is the only satellite model nematode with advanced functional genomic resources, bioinformatics and genetic tools (Rödelsperger et al. 2019). Moreover, over a quarter of humans are infected with parasitic nematodes (International Helminth Genomes Consortium 2019), which not only threaten human health, but also cause major impediment to socioeconomic development (Blaxter and Koutsovoulos 2015; GBD 2016 DALYs and HALE Collaborators 2017; Sallé et al. 2019). Parasitic nematodes also cause a variety of diseases for domesticated animals and crop plants, resulting tens of billions of dollars economic loss every year (Jones et al. 2013; Roeber et al. 2013). All the nematodes sequenced so far are either terrestrial free-living or parasitic. However, none of the large number of free-living marine nematodes have had their genomes sequenced and analysed (International Helminth Genomes Consortium 2019; Smythe et al. 2019; Weinstein et al. 2019).

Oceans cover 71% of the Earth’s surface, and represents almost 99% of available habitat, which are a key element for the existence and evolution of life (Robert 1999; Peng et al. 2020). In marine sediments, free-living nematodes abound both in numbers and in local species diversity, comprising about 80% of the abundance of meiofauna (Heip et al. 1985; Danovaro et al. 2010; Nascimento et al. 2012), they play a key role in the benthic food web and ecosystem. However, the study of marine nematodes is largely limited to taxonomy and ecology (De Meester et al. 2016; Derycke et al. 2016). Molecular mechanisms underlying their evolution, development regulation and adaptation mechanisms are rarely reported, and none of marine model nematode has been reported. The bacterivorous marine nematode *Litoditis marina* (Bastian, 1865) Sudhaus, 2011, formerly known as *Rhabditis marina* or *Pellioditis marina*, is found widely distributed in the littoral zone of coasts and estuaries of European, American, African and Asian countries, and play an important role in these marine ecosystems (Tietjen et al. 1970; Kito 1981; Houthoofd et al. 2003; Derycke et al. 2016). In addition, the embryonic cell lineage of *L. marina* has been reported by Houthoofd et al. (2003). In comparison with *C. elegans*, the overall *L. marina* lineage homology is 95.5%, whereas the fate homology is only 76.4%, and most of the differences in fate homology concern nervous, epidermal, and pharyngeal tissues, suggesting that *L. marina* will be a valuable model to study developmental evolution between marine and terrestrial nematodes.

Here, we report construction of the first highly inbred lines of *L. marina*, to the best of our knowledge, via consecutive full-sibling crosses, which might be the first in dioecious marine animals. With short generation time, the inbred lines with clear genetic background reported here make *L. marina* a promising marine model animal. Next, we successfully sequenced and assembled the genome for *L. marina*, and successfully applied CRISPR/Cas9 genome editing in this marine nematode. With comparative genomics among *L. marina* and nine terrestrial nematodes, we have identified unique genomic characters for this marine nematode’s adaptation to the marine environment. In addition, ninety-two conserved genes appear to be positively selected, which may underpin the osmotic, neuronal and epigenetic changes in the marine nematode. Therefore, our report will underpin ongoing comparative work on animal development, functional genomics, environmental adaptation, and life evolution from the ocean to land or vice versa.

## RESULTS AND DISCUSSION

### Establishment of inbred lines

To the best of our knowledge, establishment of inbred lines has not yet been achieved for any marine nematodes. Here, the original wild strain, HQ1, was isolated from intertidal sediments in Huiquan Bay, Qingdao, China. In the laboratory, it can be easily cultured at room temperature on seawater-NGM agar plates (with a salinity of 30‰), by applying *E. coli* OP50 as food resource. Worms were identified as *Litoditis marina* PmIII based on mitochondrial COI gene and nuclear 18S, 28S rDNA sequences (Supplementary Table S1).

We then generated the highly inbred lines by consecutive full-sibling crosses for 40 generations. Because of inbreeding depression, totally 3,175 crosses were performed to achieve the inbred lines construction, within about 18 months (Fig. 1A). The estimated inbreeding coefficient (*F* value) of 40th generation inbred line is 0.9998, suggesting a very clear genetic background. As we know, the emerging marine invertebrate model, the starlet sea anemone *Nematostella venctensis*, is a dioecious species with asexual reproduction, its clonal stocks could be achieved by inducing regeneration, but no inbred lines were reported for it (Stefanik et al. 2013; Russell et al. 2017). Another emerging marine model animal, the annelid worm *Platynereis dumerilii* has distinct lines of more than 10 inbred generations, however lines more than 20 inbred generations have not yet been reported (Zantke et al. 2014). According to the standard of being an inbred line with at least 20 generations’ consecutive full-sibling crosses, and an inbreeding coefficient of 0.9860 (Beck et al. 2000), our highly inbred lines reported here is the first in dioecious marine animals.

**Figure 1.**
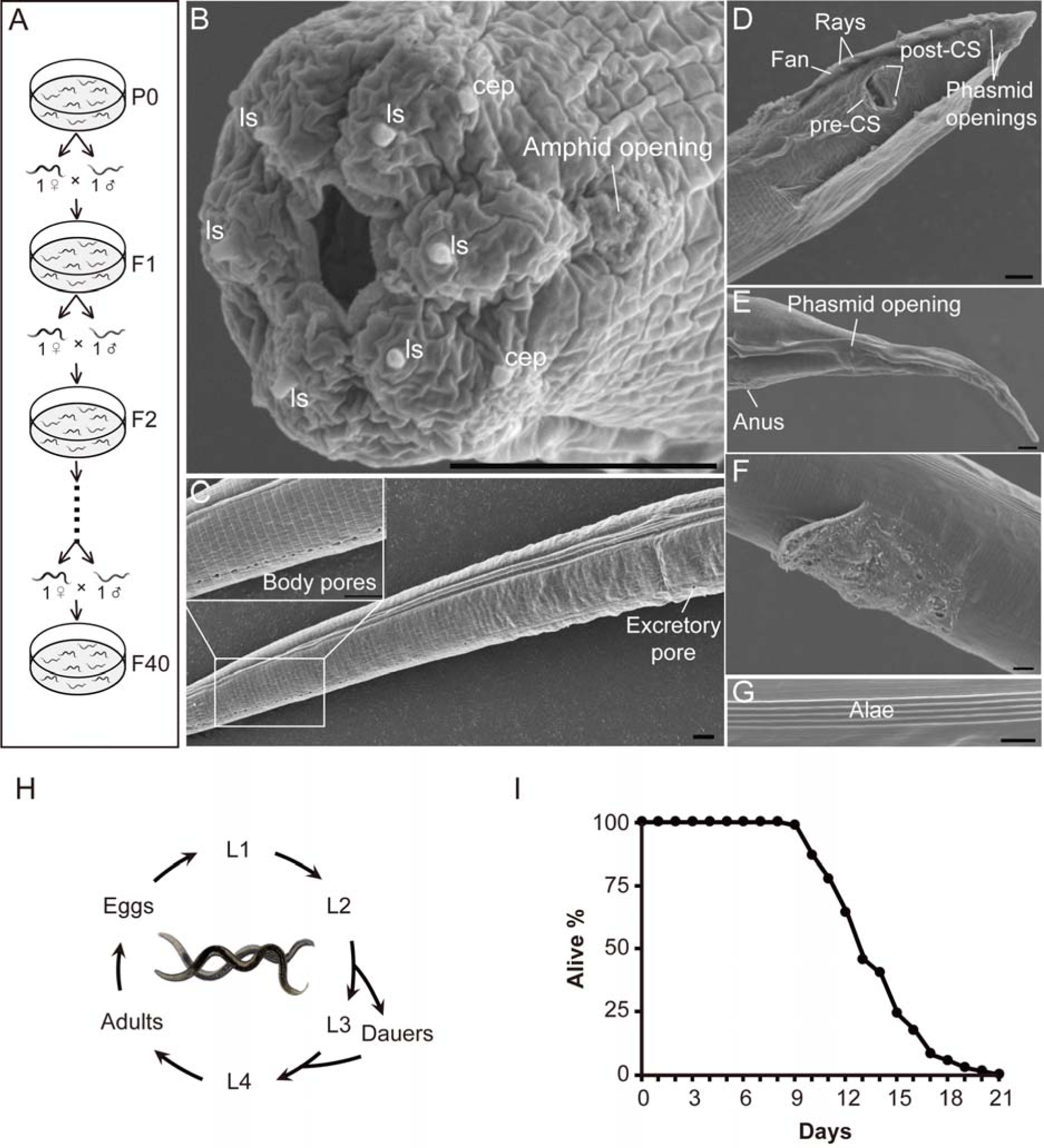
General information for *L. marina*. (*A*) Crossing scheme used to generate the inbred lines for *L. marina*. B to G Scanning electron microscopy of *L. marina*. (*B*) Mouth of an adult male. (*C*) Body pores. (*D*) Male tail. (*E*) Female tail. (*F*) Female vulva covered with plug. (*G*) Alae, cuticular lateral ridges. (*H*) Life cycle of *L. marina*. (*I*) Life span of *L. marina*. Worms in B-G are from the 23rd generation (F23) of the inbred line. cep, cephalic sensillum; CS, cloacal sensilla; ls, labial sensillum. Scale bars: 5 μm.

As an emerging marine model organism, the highly inbred line of *L. marina* poses the following experimental advantages: (a) Short generation time. At room temperature, newly hatched L1 larvae reached sexual maturation within about 4-5 days, L4 worms can live for up to 21 days till death (Fig. 1I). (b) Transparent body for imaging and large brood sizes for genetics. An average of 143 eggs (up to 369) can be produced by a single gravid female of *L. marina* (Supplementary Table S2). (c) Standard protocols for its laboratory culture and preservation. Like the model organism *C. elegans, L. marina* is well suited to rearing under laboratory conditions, and can be frozen and stored at −80 °C. (d) A homogeneous genetic background. With reduced inter-individual variation, these inbred worms will facilitate accurate design of single guide RNAs (sgRNAs) without concern for sequence polymorphisms, which is indispensable for high efficiency of CRISPR/Cas9 gene editing to modify gene expression, finally helping to uncover gene functions via large scale forward and reverse genetics. Thus, the highly inbred line of *L. marina* is an ideal system to study nematode functional genomics, developmental evolution, as well as adaptation mechanisms to sediment and global climate changes.

### *L. marina* general descriptions

*L. marina* is gonochoristic. With a transparent and cylindrical body shape, adults are 1.0-2.0 mm in length, in terms of which females slightly larger than males (Supplementary Fig. S1). The outer side of the mouth is endowed with six labial sensilla, two pocket-like amphid openings, and four cephalic sensilla (Fig. 1B; Supplementary Fig. S2), some of these morphological characters are comparable to pictures as shown by De Meester et al., although without detailed description in their report (De Meester et al. 2016). Strikingly, we observed that both female and male have protruded cephalic sensilla in *L. marina* (Supplementary Fig. S2), while only males hold for protruded cephalic sensilla in *C. elegans*. There is a pre-cloacal sensillum-like papilla and two post-cloacal sensilla around male cloacal opening (Supplementary Fig. S3A), similar to *C. elegans*. However, *L. marina* does not have a hook-like structure, and has a much contracted fan compared with *C. elegans* males, the morphology of spicules in *L. marina* is also different from that in *C. elegans* (Supplementary Fig. S3B). Furthermore, Copulatory plugs can be observed covering the vulval region of mated females (Fig. 1F), which might be associated with mate competition (Barker 1994; Palopoli et al. 2008). For both sexes, five cuticular ridges (alae) can be seen in the lateral field of adults (Fig. 1G), in contrast with three parallel ridges of alae in *C. elegans* (http://www.wormatlas.org). Interestingly, three or even more than 20 body pores were observed along the ventral and dorsal midline of *L. marina* body (Fig. 1C), while no such body pores were reported in *C. elegans*. The increased number for body pores may be a common feature for certain marine nematodes, like *Ptycholaimellus ponticus* as mentioned by Tahseen (Tahseen 2012). Besides, *L. marina* shares developmental characters with *C. elegans*, four larval stages, with an alternative dauer stage under harsh conditions (Fig. 1H). Mating experiments for *L. marina* show an obvious female-biased progeny sex ratio (Supplementary Table S2), which could be caused by mate competition (Hamilton 1967).

These substantial morphological differences between *L. marina* and *C. elegans*, particularly protruded cephalic sensilla in female *L. marina*, the loss of hook and much contracted fan as well as the different morphology of spicules in male *L. marina*, the increased number for body pores and ridges of alae in both sexes of *L. marina*, might reflect special neuronal sensation, male mating behaviour, reproduction strategies, as well as adaptation mechanisms to this marine nematode’s habitat marine environment.

### Genome assembly and phylogenetic analysis

We generated 12.05 Gb of paired-end (PE) reads (65×) with an Illumina HiSeq platform and 26.11 Gb PacBio long reads (141×) from a F23 inbred line developed by this study (Supplementary Tables S3, S4). A total assembly of 185.29 Mb, consisting of 75 contigs with an N50 contig size of 5.04 Mb, was achieved using Canu-based pipeline (Koren et al. 2017) (Supplementary Table S5). The *L. marina* genome assembly showed high integrity and quality. Of total Illumina reads, 93.93% could be mapped to the assembly. According to CEGMA analysis (Parra et al. 2007), 453 out of 458 core eukaryotic genes are present in the assembly. The completeness of gene regions was further assessed using BUSCO (Simão et al. 2015), which showed that 92.36% of the Nematoda single-copy orthologs are complete in the assembly (Supplementary Table S6).

The annotation for *L. marina* genome was built on protein homology-based prediction, RNA-Sequencing-based approach and *de novo* prediction, which contains 17,661 protein-coding genes (Table 1; Supplementary Table S7). Compared to the terrestrial free-living nematode *C. elegans, C. nigoni* and *P. pacificus, L. marina* has a notable larger genome size (Table 1). In *L. marina*, we identified 30.4% of the assembly as repetitive elements, which is much higher than that reported for *C. elegans* and *P. pacificus* (Table 1). With about 56.33 Mb repetitive elements in *L. marina* compared to 18.3 Mb in *C. elegans*, the expansion of repeats certainly contributes a significant part to its genome size expansion. Our data confirmed that repetitive sequences take up much lower ratio of genome in selfing species than outcrossing species (Table 1), regardless of genome sizes, which might infer repeat shrinkage during selfing process. In addition, we found that *L. marina*’s average intron size is significantly larger (5,353 bp), resulting in a striking larger average gene size (7,004 bp), compared to *C. elegans* (3,143 bp) and other terrestrial free-living nematodes (Table 1). Furthermore, average exon number per gene was estimated as 12 in *L. marina*, which is similar to that in *P. pacificus*, but much higher than that in both *C. elegans* and *C. nigoni*. Additionally, we found that many of single contigs in *L. marina* showed synteny with more than one chromosome of *C. elegans* and *C. nigoni*, suggesting striking genomic evolutionary histories (Fig. 2A; Supplementary Fig. S4).

**Table 1.** Genome statistics.

| Table 1 Genome statistics |  |  |  |  |
| --- | --- | --- | --- | --- |
|  | *L. marina* | *P. pacificus* | *C. elegans* | *C. nigoni* |
| Estimated genome size (Mb) | 178.1 | 161.7 | 100 | 129.5 |
| Final assembly size (Mb) | 185.3 | 158.5 | 100.3 | 129.5 |
| Contig N50 (bp) | 5,044,337 | 4,619,064 | 17,493,829 | 3,254,670 |
| Contig N90 (bp) | 1,737,331 | 812,026 | 13,783,801 | 562,841 |
| GC content of whole genome (%) | 36.1 | 42.8 | 35.4 | 37.8 |
| Repetitive sequence (%) | 30.4 | 22.4 | 18.3 | 29.9 |
| Content of transposable element (%) | 23.3 | 17.9 | 16.6 | 23.9 |
| Content of SSR (%) | 0.05 | 0.04 | 0.29 | 2.8 |
| Proportion of genome that is coding (%) | 66.8 | 73.7 | 63.2 | 49.1 |
| Number of putative coding genes | 17,661 | 25,517 | 20,178 | 29,167 |
| Average gene size (bp) | 7,004 | 4,579 | 3,143 | 2,181 |
| Average CDS size (bp) | 1,328 | 1,350 | 1,233 | 1,142 |
| Average intron size (bp) | 5,353 | 3,213 | 1,592 | 1,022 |
| Average CDS number per gene | 12 | 13 | 6 | 5 |
| GC content in coding region (%) | 36.7 | 43 | 36.4 | 39.6 |

**Figure 2.**
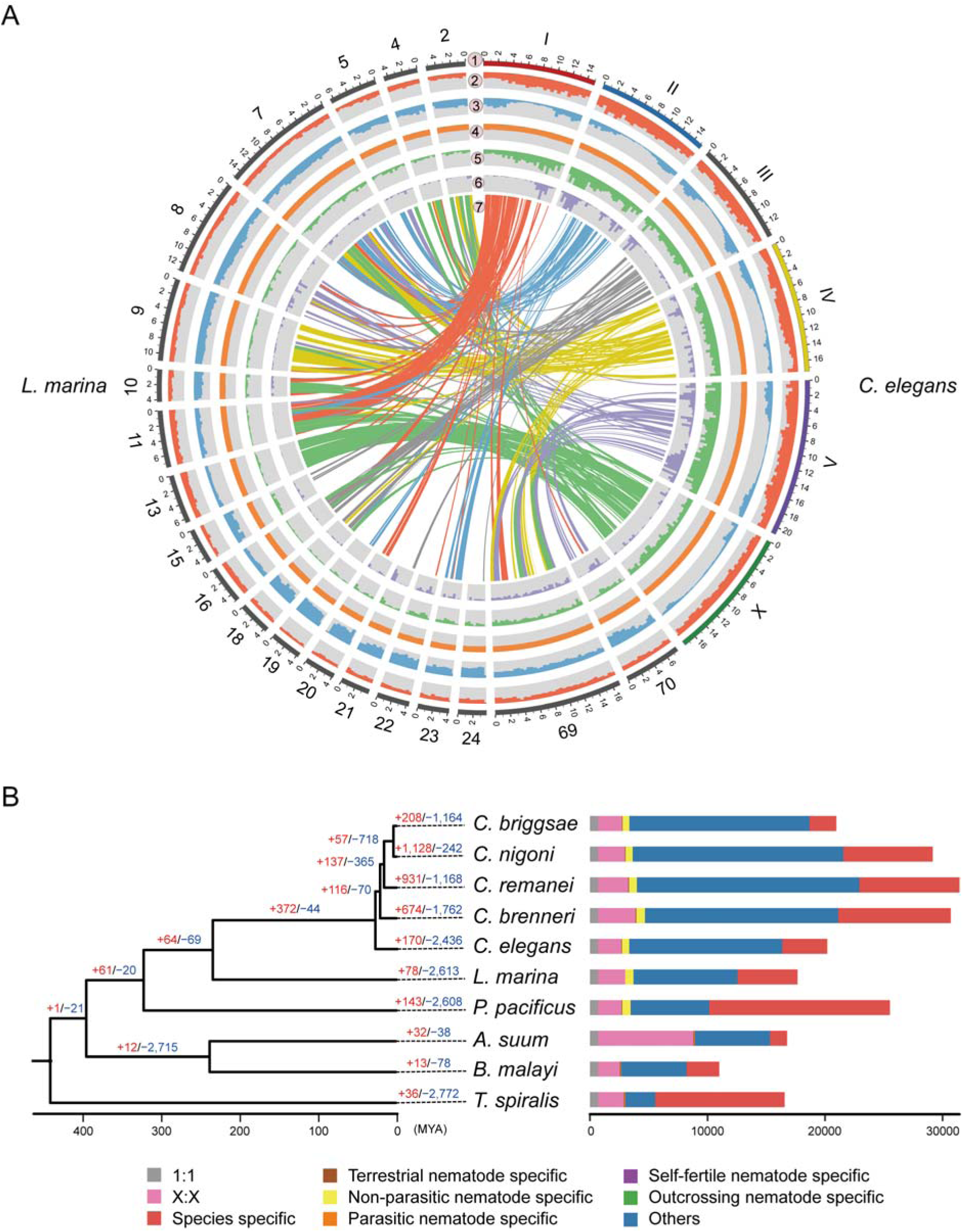
Phylogenetic and evolutionary comparisons between *L. marina* and terrestrial nematodes. (*A*) Comparative analysis of genomic features of *L. marina* vs *C. elegans* over 500 Kb chromosomal intervals. Tracks from outside to inside: 1, positions (in Mb) of the six chromosomes of *C. elegans* and Top 20 contigs in length for *L. marina*; 2, distribution of gene density; 3, repetitive DNA density; 4, GC content; 5, unique genes density; 6, distribution of expanded gene families; 7, DNA sequence synteny. (*B*) Phylogeny assignment of 10 species. Numbers labelled on each branch are the specific number of gene gain (+) or loss (-). The estimated divergence times are displayed below the phylogenetic tree. Orthologous genes are categorized in stack bar, and the length of stack bar is proportional to number of genes. The 1:1 set denotes single-copy orthologs that were present in all species. X:X indicates orthologs present in multiple copies in individual species. Species specific genes were those that had no homologs in the other species used in this analysis.

Applying OrthoMCL to *L. marina* and nine terrestrial nematodes, including both free-living (*C. elegans, C. nigoni, C. briggsae, C. brenneri, C. remanei, P. pacificus*) and parasitic (*A. suum, B. malayi, T. spiralis*) species, we identified a total of 27,134 orthogroups. Among these orthogroups, 697 contained putative single-copy gene families. Phylogenetic trees were constructed with the maximum likelihood method (Stamatakis 2014) (Fig. 2B). The divergence time estimation indicated that *L. marina* and the genus *Caenorhabditis* have diverged approximately 234 million years ago, and the divergence time between *L. marina* and *P. pacificus* was estimated to be about 88 million years (Fig. 2B). As *L. marina* and *Caenorhabdits* worms belonging to the same family, together with a *C. elegans* satellite model nematode *P. pacificus*, they belong to the same order. Thus, this marine nematode and its terrestrial relatives provide a comparative research platform to study how an animal adapts to its habitat environment (ocean or land), and the mechanism underlying life evolution from the ocean to land, or vice versa.

### CRISPR/Cas9 genome editing in *L. marina*

To facilitate *L. marina* to be a model animal, we applied CRISPR/Cas9 gene editing for further gene function studies. We performed knockout experiments using CAS9 protein and in vitro synthesized single guide RNA (sgRNA) targeting the second exon of *Lma-rol-6*. Excitingly, we identified a homozygous roller mutant line with an 8 bp deletion in *Lma-rol-6* gene around the sgRNA target site (Fig. 3). As far as we know, this is the first successfully application of gene editing in marine nematodes, which will facilitate our ability to further study the functions of interesting genes in *L. marina*.

**Figure 3.**
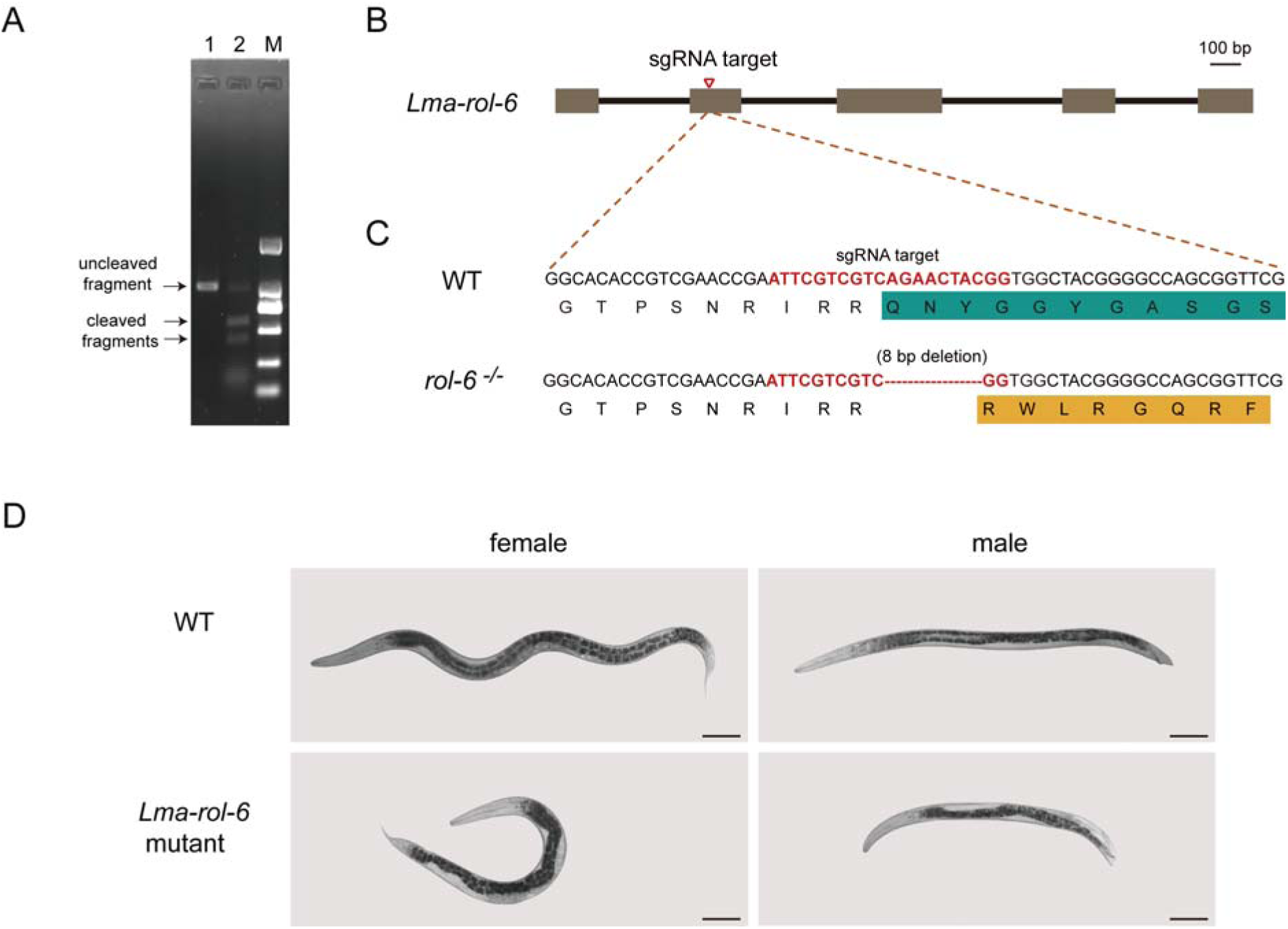
*rol-6* CRISPR/Cas9 genome editing in *L. marina*. (*A*) In vitro cleavage assay. Most *Lma-rol-6* target fragment (992 bp) was cleaved into two fragments of 568 bp and 424 bp by RNPs. Lane 1 shows the untreated *Lma-rol-6* target fragment, lane 2 shows the treated *Lma-rol-6* target fragments. Lane M shows DL2,000 DNA Marker (Takara, 3427A), the bands from bottom to top are 100 bp, 250 bp, 500 bp, 750 bp, 1,000 bp and 2,000 bp. (*B*) Schematized *Lma-rol-6* gene highlighting the sgRNA target site in the second exon. Scale bars: 100 bp. (*C*) Mutation sites identified in *Lma-rol-6* mutant line by sequencing. (*D*) Comparison of wild type (WT) worms and representative *Lma-rol-6* mutants. Scale bars: 100 um.

### Transposable elements in *L. marina*

In *L. marina*, TEs occupied 23.3% of the genome assembly length, which accounts about 43.3 Mb (Table 1). To determine the detail composition of TEs, we further analysed them in *L. marina* and *C. elegans* based on methods reported by Szitenberg et al. (2016), and conducted comparative study with other six terrestrial nematodes. We found that *L. marina* resembles *P. pacificus* most in TEs composition (Supplementary Table S8). For class I TEs, the number increases in *L. marina* compared to other nematodes except *P. pacificus* (Supplementary Table S8). For class II DNA transposons of *L. marina*, there is a decrease both in the number and percentage compared to those of *Caenorhabditis* species. DNA transposon families, e.g., EnSpm, TcMar, Helitron and hAT, contribute largely to this contraction (Supplementary Table S8).

Compared to *C. elegans, L. marina* has increased numbers of TEs in long terminal repeat (LTR) retrotransposons. In this article, we defined intact LTR retrotransposon as repeat sequence bounded by LTRs, having a primer binding site (PBS), a polypurine tract (PPT), as well as LTR retrotransposon domains including protease, reverse transcriptase, RNase H, and integrase. We found 1 intact LTR retrotransposon in *C. elegans*, 9 in *C. inopinata*, the sibling species of *C. elegans* with expanded LTRs, and 10 in *L. marina* using combination of LTRharvest and LTRdigest (Supplementary Table S9). Partial LTR retrotransposon is defined as repeat sequence bounded by LTRs with at least one of the domains mentioned above but having both PBS and PPT. We found 5 partial LTR retrotransposons in *C. elegans*, 16 in *C. inopinata* and 39 in *L. marina* (Supplementary Table S9). In addition, homolog-searching method resulted in 22 LTR retrotransposons in *C. elegans*, 119 in *C. inopinata* and 91 in *L. marina* (cut off E-value =10-7; Supplementary Table S9). By LTR classification criteria of Kanzaki et al. (2018), we found 5 full LTR retrotransposons and 17 partial LTR retrotransposons in *C. elegans*, 65 full LTRs and 125 partial LTRs in *C. inopinata*, 51 full LTRs and 123 partial LTRs in *L. marina* (Supplementary Table S9). Together, these data show that LTR retrotransposons in *L. marina* and *C. inopinata* expand significantly, compared to the model organism *C. elegans*.

The ecological niche occupied by *P. pacificus* is completely different from that of *C. elegans. Pristionchus* nematodes live in close association with beetles in a nearly species-specific manner (Dieterich et al. 2008). Similarly, although known habitats of most other *Caenorhabditis* species are soil, rotting organic materials, and leaf-litter environments, *C. inopinata* was isolated from fresh syconia of the fig *Ficus septica* and is likely to have a close phoretic association with the fig wasp *Ceratosolen bisulcatus* (Kanzaki et al. 2018). We thus wonder that the TEs’ similarity between *L. marina* and *P. pacificus*, the LTR expansions in both *L. marina* and *C. inopinata*, might suggest not only their phylogenetic similarity, but also certain similar evolutionary selection in their life histories. Systematic detection of such differences and understanding of their roles in vivo remains a significant challenge.

### Nuclear hormone receptors in *L. marina*

Based on results of gene family and protein domain analysis, we found appreciable contractions in nuclear hormone receptor (NHR) gene families in *L. marina* compared to other free-living nematodes. 96 putative NHRs were identified in *L. marina* by manual curation, while the number of NHRs varies within terrestrial nematode species reaching 275 in *C. elegans*, 166 in *C. briggsae* and 272 in *C. remanei* (Fig. 4A).

**Figure 4.**
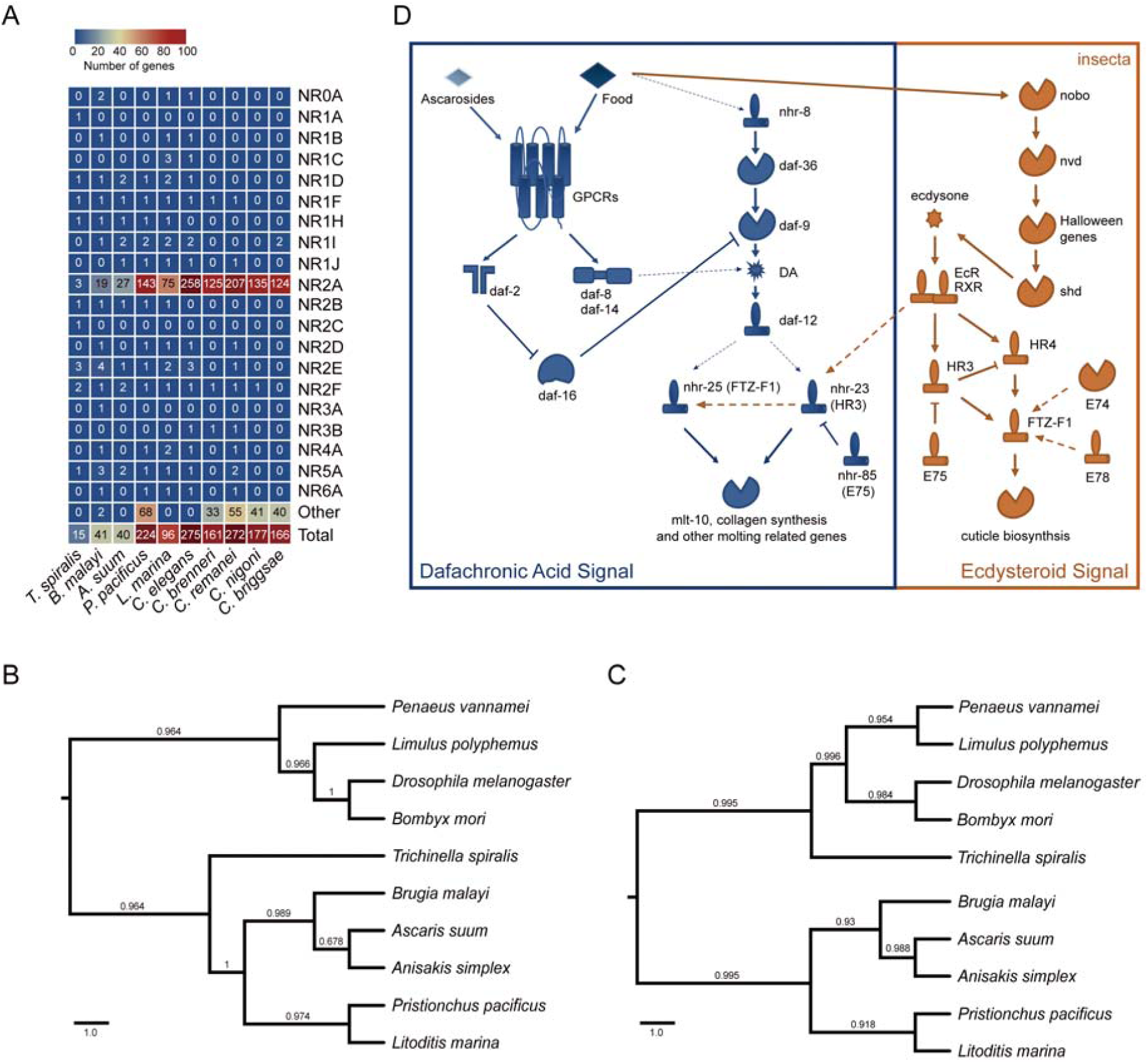
Diversity of NHRs in *L. marina*. (*A*) A heat map showing gene numbers of NHRs in *L. marina* and comparison with other nematodes. (*B*) Phylogenetic analysis of EcRs. Putative EcRs in *L. marina* (EVM0014194.1) and *A. simplex* (ASIM_0001371101-mRNA-1) were identified based on BLASTP and hmmscan, EcRs of *A. suum* (ADY42041.1), *B. malayi* (ABQ28713.1), *B. mori* (XP_021206379.1), *D. melanogaster* (NP_724456.1), *L. Polyphemus* (XP_022239580.1), *P. pacificus* (ACY82385.1), *P. vannamei* (ROT69959.1) and *T. spiralis* (XP_003376657.1) were downloaded from NCBI with indicated Genebank number. Values on branches are genetic distance. (*C*) Phylogenetic analysis of RXRs. One putative RXR in *L. marina* (EVM0012017.1) was identified based on BLASTP. RXRs of *A. simplex* (ASIM_0001060801-mRNA-1), *A. suum* (AgB07_g019_t01), *B. malayi* (Bm3989.1), *B. mori* (XP_012548309.1), *D. melanogaster* (NP_476781.1), *L. Polyphemus* (XP_022251212.1), *P. pacificus* (ACY82384.1), *P. vannamei* (AGG55291.1) and *T. spiralis* (EFV60819) were downloaded from NCBI. Values on branches are genetic distance. (*D*) Model of putative EcR and RXR function in *L. marina*. We proposed that EcR and RXR work coordinately with DAF-12 in regulating development and molting in *L. marina*. Dafachronic acid signaling, marked by blue, is reported in *C. elegans*, while the ecdysteroid signaling (in orange) represents *D. melanogaster* orientation. Environmental signals, e.g., food and ascarosides, are transduced through the TGF-β and insulin pathway regulating both DAF-12 and EcR/RXR. Interactions among these NHRs, homologs of early-late genes, governate molting cycles and larval transitions. All the processes in this diagram do not commence simultaneously. DA, dafachronic acid.

Phylogenetic analysis suggested that the contraction of *L. marina* NHRs mainly occurred in groups which are inferred as supplementary nuclear receptor genes on *C. elegans* chromosome V (Robinson-Rechavi et al. 2005) (Supplementary Fig. S5). NHRs such as dpr-1 (NR3B), sex-1 (NR1D), nhr-8 (NR1J) and nhr-114 (NR2A), related to resistance, reproduction, sex-determination and developmental regulation, were absent in *L. marina*. These contractions suggested that rather stable living environment of *L. marina* might require less NHRs.

Homologs of evolutionarily conserved NHRs acting as major regulators were identified in *L. marina*, include homologs of PPAR (NR1C), Rev-Erb (NR1D), ROR (NR1F), COUP (NR2F), and NGF-1 (NR4A1) (Antebi 2015). Expansion of several NHRs was also observed in *L. marina* but most of their homologs’ function in *C. elegans* remains unclear. For example, 5 homologs of *C. elegans* hypothetical protein Y67D8B.2 and 8 homologs of NHR-173 were present in *L. marina*, indicating divergence of NHRs in these two nematodes. Strikingly, putative ecdysone receptor (EcR) and retinoic X receptor (RXR), which are absent in *C. elegans* and most free-living nematodes (Schumann et al. 2018), were found in *L. marina* (further confirmed by both RNA-Seq data and genomic PCR). Ecdysteroid signaling was widely observed in filarial parasitic nematodes and putative homologs of RXR and EcR were also identified in beetle-associated nematode *P. pacificus* and its relative species (Parihar et al. 2010; Kostrouchova and Kostrouch 2015; Prabh et al. 2018). Homologs of *D. melanogaster* EcR and RXR pathway genes HR3 (NHR-23), FTZ-F1 (NHR-25) and E75(NHR-85) (Fig. 4D) were all identified in *L. marina*, indicating the existence of ecdysteroid signaling in *L. marina*. Phylogenetic trees of EcRs and RXRs also reveals the high similarity between *L. marina* and *P. pacificus* (Fig. 4B-C). Since the necromenic lifestyle in *P. pacificus* shares high similarity with parasitism, we propose that there might be an exogenous ligand regulating *L. marina* development, morphology and behavior via animal-animal communications, similar to that in beetle and *P. pacificus* pair’s relationship (Werner et al. 2017).

In Ecdysozoa, molting is a major developmental timing landmark in life cycle. The endocrine and molecular control of molting have been well characterized in insects (Hada et al. 2010). Ecdysone, whose levels oscillates during the larvae development, acts through a heterodimeric receptor composed of EcR and RXR/Usp, to activate genes responsible for physiological changes leading to larval ecdysis, including HR3, E74, E75, E78 and FTZ-F1 (Riddiford et al. 2003; Kostrouchova and Kostrouch 2015; Prabh et al. 2018) (Fig. 4D). Although numerous studies have been conducted on *C. elegans* and other nematodes, the mechanism and regulation of molting remains obscure. Exogenous ecdysteroids showed effects on molting and reproduction especially for filarial parasites (Barker et al. 1990; Tzertzinis et al. 2010), and EcR and RXR homologs in *B. malayi* have been proved to be functional, which can respond to 20-hydroxyecdysone, an active form of ecdysone in most insects (Tzertzinis et al. 2010). In addition, genes encoding the putative RXR/Usp and EcR homologs in *P. pacificus*, has been shown to have cyclical peaks of expression prior to all three post-hatching molts, indicating the possible role in molting regulation (Parihar et al. 2010; Lewis and Hong 2014). In *C. elegans*, two nuclear hormone receptors, NHR-23 (HR3) and NHR-25 (FTZ-F1), oscillating in parallel with the molting cycle and synthesis of cuticular collagens, regulates its molting (Hada et al. 2010). In addition, it’s been reported that the highly conserved nuclear hormone receptor DAF-12, which is related to the vitamin D receptor, indirectly regulates NHR-23 and NHR-25 levels in *C. elegans* (Patel and Frand 2018). Both DAF-12 and EcR/RXR pathway genes were observed in *L. marina* genome. Collectively, we infer that EcR and RXR may work coordinately with DAF-12 in regulating development and molting in *L. marina*. Furthermore, as EcR and RXR being reported in parasitic nematodes as major players in molting and development, this free-living nematode might be a useful model animal for research on pathogenic nematodes, and developing new drugs for parasitic nematode diseases in crops, living stocks and human beings.

### Expansion of immunity and oxygen regulation genes

The marine environment, consisting of the open ocean, the upper 10-50 cm of the ocean seafloor sediment, as well as the underlying oceanic deep subsurface, is primarily occupied by microbes, mainly bacteria and protists, which account for about 70% of the total marine biomass (Bar-On et al. 2018). In addition, bacteria in marine sediments are more phylogenetically diverse than in any other environments (Lozupone and Knight 2007). Thus, the benthic marine nematode *L. marina* must yield strategies to deal with not only the relatively low oxygen level (Breitburg et al. 2018), but also putative more complex bacterial communities in its habitat.

The most significant expansion of immunity related genes was found in C-type lectin domain proteins (Fig. 5A). Genes encoding DAF-19 and p53 are also expanded compared to the terrestrial nematodes (Fig. 5A). DAF-19, an ortholog of regulatory factor X (RFX) transcription factor, which was initially discovered in human immune cells, has conserved roles both in cilia formation and immunity across different phyla (Reith and Mach 2001; Piasecki et al. 2010). In *C. elegans*, the sole *daf-19* gene has been reported regulating innate immunity in response to pathogenic bacterial food (Xie et al. 2013). Moreover, roles in sensing or responding to oxygen of DAF-19 and p53 have been shown in *P. pacificus* (Moreno et al. 2018) and *C. elegans* (Derry et al. 2001), respectively. IAdditionally, carbohydrate sulfotransferase genes, involved in diverse extracellular recognition processes (Bowman and Bertozzi 1999), are also notably expanded in the *L. marina* genome (Fig. 5A). Taken together, these features might facilitate enhancing *L. marina*’s adaptation to habitat oxygen levels, and surrounding bacterial environment.

**Figure 5.**
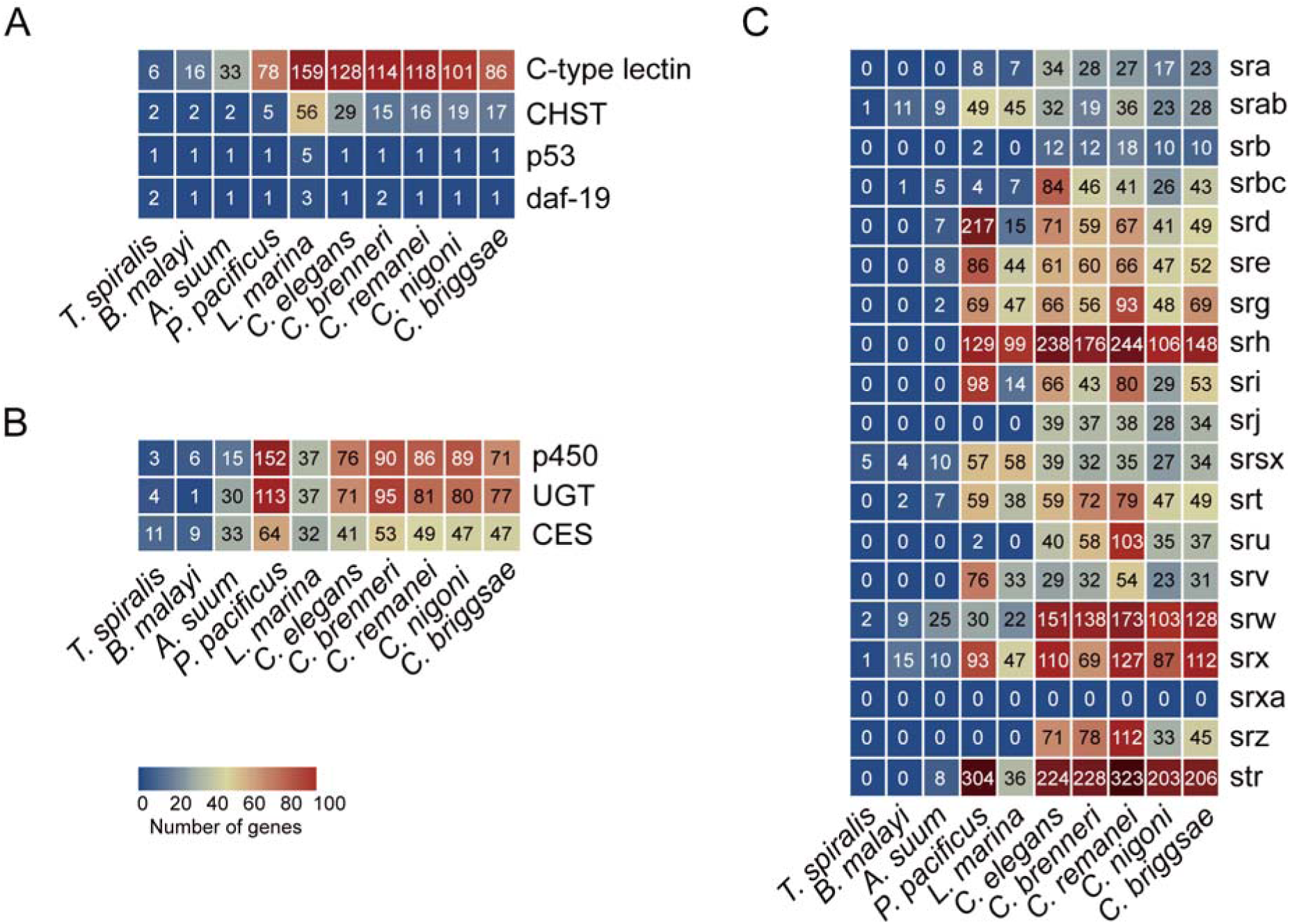
Gene or gene family expansions and contractions. (*A*) Gene expansions in *L. marina*. CHST, carbohydrate sulfotransferase. (*B*) Gene contractions in both *L. marina* and parasitic nematodes. CES, carboxylesterase; UGT, UDP-glucuronosyltransferase. (*C*) Diversity of serpentine receptors in different nematodes.

### Gene family contractions in xenobiotics detoxification, serpentine receptors and histones

Interestingly, we found that several gene families, such as cytochrome P450, UDP-glucuronosyltransferases (UGTs) and carboxylesterases (CESs), which play crucial roles in the detoxification of xenobiotics (Lindblom and Dodd 2006; Xu et al. 2016), are contracted greatly in *L. marina* compared to its land relatives (Fig. 5B). Additionally, we also observed notable contractions in seven-transmembrane chemoreceptor gene families in *L. marina* (Fig. 5C), especially for srb, srj, sru and srz serpentine families. Gene families related to sra, srbc and srw all showed shrinkage in *L. marina, P. pacificus* and the tested parasitic species. While 6 other families (srd, srh, sri, srt, srx and str) were only contracted in *L. marina* and parasitic ones. These results suggest that *L. marina* requires a clearly narrow range of chemoreceptors to detect possibly only a small variation of chemical stimuli in its habitat area, as the case for the parasitic nematodes. Collectively, we thus inferred that these contractions in *L. marina* contributes to a relatively stable marine environment compared to the terrestrial scenarios.

In eukaryotes, histone proteins are key players for DNA package and gene regulation. There are five types of histone genes, coding for the linker histone H1 and four core histones, H2A, H2B, H3 and H4. These histones are typically classified as two major categories, one is the canonical replication-dependent histones, and the other is the histone variants, isolated as single-copy genes with introns and are typically constitutively expressed (Marzluff and Koreski 2017). Base on genome annotation, we identified totally 43 histone genes in *L. marina*, including 10 H1 genes and 33 core histone genes. It revealed that the number of all core histone genes, especially for H4 and H2B, is apparently contracted compared with *C. elegans* and other *Caenorhabditis* worms (Fig. 6A; Supplementary Fig. S6), indicating a decreased diversity for histones in *L. marina*. Furthermore, we found that the portion of core histone variants is significantly higher in *L. marina* (33%) than that in *C. elegans* (11%). On the other hand, for canonical histones, the majority of that in *C. elegans* are identified as single-copy genes, while a considerable number of that in *L. marina* containing multiple copies. Therefore, these features for histone genes could be correlated with its adaptation to the more defined marine environment for *L. marina* as well.

**Figure 6.**
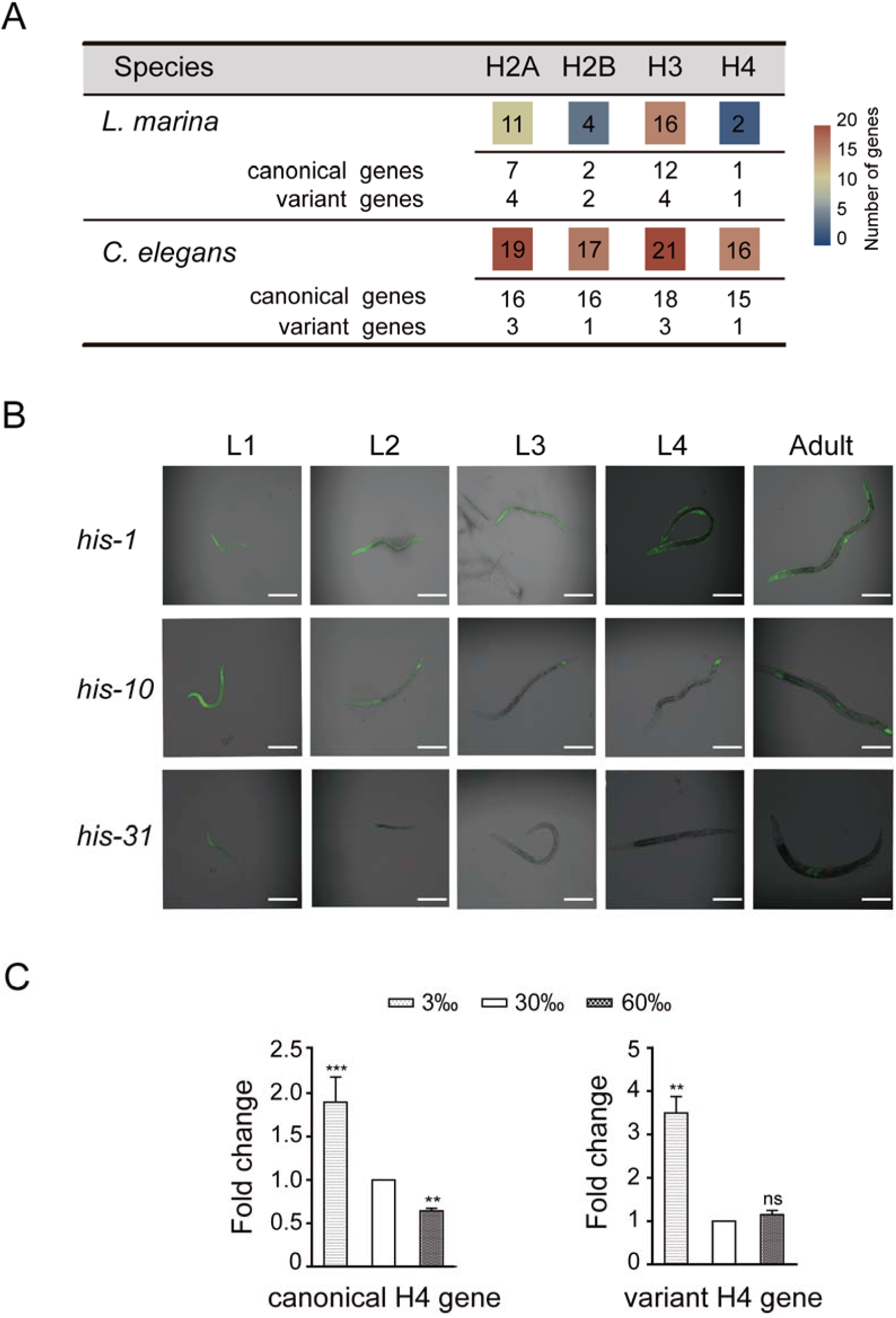
Core histones contractions in *L. marina*. (*A*) Gene numbers for four types of core histones between *L. marina* and *C. elegans*. (*B*) Expression patterns for three H4 genes in *C. elegans*. Scale bars: 150 μm. (*C*) qPCR analysis of two *L. marina* H4 genes under different salinity conditions (3‰, 30‰, 60‰).

Notably, there are only two H4 genes in the marine nematode, one is a canonical H4 with about 57 copies in the genome and the other one is a single H4 variant gene. By contrast, 16 H4 genes exist in *C. elegans*, which are classified into 5 groups based on similarity analysis on their promoter sequences (Supplementary material). Using transgenic worms, we further investigated expression patterns for 3 different types of *C. elegans* H4 genes, *his-1, his-10* and *his-31*, respectively. In spite of the fact that 3 genes all showed highest expression levels in L1 worms and embryos, they did demonstrate different spatial expression patterns as shown in Figure 6B, suggesting a mechanism to cope with complex terrestrial environments for *C. elegans*. To ask whether *L. marina* H4 genes are involved in responding to environmental variations, we performed qPCR for both H4s under different salinity conditions. The results exhibited that expression levels are significantly elevated for both genes under lower salinity condition, while it was decreased obviously for the canonical H4, but not the H4 variant, under higher salinity (Fig. 6C), indicating two *L. marina* H4 genes functioning differently to react the same environmental perturbations. Furthermore, unlike the case of 17 H2B genes randomly distributed on different chromosomes in *C. elegans*, we found only 4 H2B genes in the *L. marina* genome, one of which is a H2B variant located adjacent to the H4 variant mentioned above. Therefore, we proposed that certain coordinately regulation mechanism might exist among the limited divergent histone genes in *L. marina*, which could be correlated with its habitat adaptation.

Additionally, based on protein domain analysis among 10 selected nematode species in our paper, nearly a hundred of protein families are contracted in both marine nematode and parasitic ones, compared to the terrestrial free-living nematodes (Supplementary Table S10), suggesting that marine environments are in large defined and more stable than the land. Thus, as marine animals already adapt to a relatively more stable environment, it is possible the reason why certain marine animals are more sensitive to the global climate change compared to the terrestrial ones (Pinsky et al. 2019).

### Positively selected genes in *L. marina*

The successful inhabiting in marine environments demands animal’s adaptation to the much higher osmotic conditions (around 30‰ in salinity) compared to the terrestrial living systems. To counter this osmotic stress, the accumulation of organic osmolytes to compensate for water loss is a universal strategy of cells (Yancey et al. 1982). Glycerol and trehalose are known osmolytes used in nematode (Lamitina and Strange 2005; Lamitina et al. 2006; Tahseen 2012; Choe 2013). Glycogen acts as a reservoir not only for energy supply, but also for osmolytes production (Frazier and Roth 2009; Possik et al. 2015). We observed an increased number of genes related to glycogen biosynthesis (Supplementary Fig. S7). There were 4 genes encoding glycogen synthases in *L. marina* versus 1 in *C. elegans*. Moreover, the strong positive selections on EVM0011198.1(*col-109*), EVM0009368.1(*lec-1*), EVM0011012.1 (*T13H5*.*6*), EVM0014994.2(*osm-12*), EVM0011797.1(*gem-1*), and EVM0008924.1(*imp-2*) homologue genes in the sea nematode genome might indicate an adaptation in osmoregulation necessary for marine life. *col-109* is predicted to be a structural constituent of cuticle, as we know the cuticle is the first layer of osmotic defence barrier and an osmotic stress damage sensor (Dodd et al. 2018). *lec-1* is mainly localized in the collagen and cuticulin-based cuticle extracellular matrix, and plays a defensive role against damage due to oxidative stress (Takeuchi et al. 2013). *T13H5*.*6*, a human Na, K-ATPase Interacting protein 4 homologue, regulates sodium ion transport, which might be essential to osmotic regulation (GO:0002028). *osm-12*, an ortholog of human BBS7 (Bardet-Biedl syndrome 7) is involved in chemotaxis and cilium assembly, *osm-12(n1606)* mutant worms showed significantly higher survival at 400 and 500 mM NaCl (Lee et al. 2016). *gem-1*, an ortholog of human SLC16A14 (solute carrier family 16 member 14) and SLC16A9 (solute carrier family 16 member 9), is predicted to be involved in monocarboxylic acid transport (GO:0015718). *imp-2* is predicted to be related in ecdysis, collagen and cuticulin-based cuticle related functions (GO:0042395). Taken together, the results indicate that the marine nematode has changed its genetic base to adapt the higher salinities in the sea. Functional experimental evidence can test how these changes affect the osmotic regulatory system.

It has been reported that the overall cell lineage homology between *L. marina* and *C. elegans* is 95.5%, while the fate homology is only 76.4%, most of the differences concern nervous, epidermal, and pharyngeal tissues (Houthoofd et al. 2003). However, the genetic bases underlying the differences are unknown. Notably, genes, that are functioning in neuronal differentiation, development and behaviour regulation, such as EVM0013824.1(*sox-2*), EVM0014071.1(*fax-1*), EVM0016776.1(*cab-1*), EVM0004317.1(*itsn-1*), EVM0003699.2(*ceh-14*), EVM0005599.1(*ceh-16*), EVM0002661.1(*ldb-1*) and EVM0006320.1(*tax-4*), were identified under positive selection in *L. marina. sox-2*, an HMG-box transcription factor, regulates olfactory neuron differentiation in *C. elegans* (Alqadah et al. 2015). *fax-1*, a nuclear hormone receptor regulates axon pathfinding and neurotransmitter expression (Much et al. 2000). *cab-1* is an ortholog of human NPDC1 (neural proliferation differentiation and control 1), which is involved in chemical synaptic transmission, positive regulation of anterior/posterior axon guidance, and ventral nerve cord development (Iwasaki and Toyonaga 2000). *itsn-1* controls vesicle recycling at the neuromuscular junction (Wang et al. 2008). *ceh-14*, an ortholog of human LHX3 (LIM homeobox 3) and LHX4 (LIM homeobox 4), is required for phasmid function and neurite outgrowth (Kagoshima et al. 2013). *ceh-16*, an ortholog of human EN2 (engrailed homeobox 2), controls neuron differentiation (GO:0030182). *ldb-1* is necessary for several neuronal functions mediated by LIM-HD proteins, including the transcriptional activation of *mec-2*, the mechanosensory neuron-specific stomatin (Cassata et al. 2000). *tax-4*, an ortholog of human CNGA2 (cyclic nucleotide gated channel subunit alpha 2) and CNGA3 (cyclic nucleotide gated channel subunit alpha 3), regulates thermosensory neuron gene expression and function in *C. elegans* (Satterlee et al. 2004). Therefore, we supposed that these positively selected genes might answer how the fate homology deference between *L. marina* and *C. elegans* occurred, and the understudied physiology and behaviours in marine nematodes.

In addition, we found that several epigenetic regulation genes such as EVM0014609.1(*utx-1*),EVM0007128.1(*dcr-1*),EVM0011449.1(*hda-4*), EVM0006140.1(*damt-1*), EVM0005066.1(*C53A5*.*17*), were positively selected in marine nematode *L. marina. utx-1*, an ortholog of human KDM6A (lysine demethylase 6A), regulates the memory of environmental chemical exposure in *C. elegans* (Camacho et al. 2018). *dcr-1*, an ortholog of human DICER1, play major roles in production of siRNA involved in RNA interference and primary miRNA processing (Grishok et al. 2001). *hda-4*, an ortholog of human HDAC4 (histone deacetylase 4) and HDAC5 (histone deacetylase 5), regulates chemoreceptor gene expression and sensory behaviours in *C. elegans* (van der Linden et al. 2008). *damt-1*, an orthologue of human METTL4 (methyltransferase like 4), regulate 6mA levels and crosstalk between methylations of histone H3K4 and adenines and control the epigenetic inheritance of phenotypes associated with the loss of the H3K4me2 demethylase *spr-5* (Greer et al. 2015). *C53A5*.*17*, an ortholog of human TRMT5 (tRNA methyltransferase 5), is predicted to have tRNA (guanine-N1-)-methyltransferase activity. Systematic detection of such epigenetic differences and understanding of their roles might open a significant area in environmental adaptation and evolution.

## CONCLUSION

To the best of our knowledge, this is the first free-living marine nematode genome ever reported, and our inbred lines published here are the first in the marine nematode community. Together with our first successfully applying CRISPR/Cas9 genome editing in marine nematode in this study, we believe that this report will bring *L. marina* to be an important marine invertebrate model to the whole marine life science field, and a unique satellite marine experimental animal to the well-known biomedical model nematode *C. elegans*. Our report indicates that marine nematode’s genome compositions already adapt to a relatively more defined environment, with massive contractions in NHRs, P450s, chemoreceptors and core histone genes. It is possible the reason why certain marine animals are more sensitive to the global climate change compared to the terrestrial ones, which should be highly alarmed to the public, ocean administration and utilization. In addition, ninety-two conserved genes appear to be positively selected in *L. marina*, and these may underpin its osmotic, neuronal and epigenetic changes in the sea. Therefore, our *L. marina* genome and other functional genomic data, inbred lines with clear genetic background and genetic manipulation system will be essential resources for animal development, functional genomics, as well as for studying environmental adaptation, global climate change and life evolution from the ocean to land or vice versa.

## METHODS

### Nematode culture

The original *Litoditis marina* wild strain, HQ1, was isolated from intertidal sediments in October, 2016 in Huiquan Bay, Qingdao, China. Briefly, sediment samples were collected and settled on 90 mm seawater-nematode growth medium (SW-NGM) agar plates (instead of H_2_O and NaCl components for normal NGM, SW-NGM plates were prepared with seawater in 30‰ salinity), kept at room temperature (20-25 °C) in the laboratory. The propagated worms were checked regularly under an Olympus SZX7 Stereomicroscope and transferred to SW-NGM plates seeded with *E. coli* OP50.

### Establishment of inbred lines for *L. marina*

We conducted a series of consecutive full-sibling crosses to produce marine nematodes with different inbreeding coefficients as illustrated in Figure 1A. All crosses were performed on 35 mm SW-NGM agar plates, seeded with OP50. Within each generation, about 100 replicate crosses were set up. Each cross was initiated by mating a single randomly selected L4 male to a randomly selected virgin L4 female. Always keep 3-10 plates with abundant offspring for next cross. In total, 3,175 crosses were set up till the final 40th generation. The 23rd generation inbred line (F23) of *L. marina* was used in the following study.

### Scanning electron microscopy

Male and female adults were individually selected and washed in filtered seawater, then fixed with 2.5% glutaraldehyde for 2 h at room temperature. After fixation, worms were rinsed twice in sterilized water and dehydrated through an ethanol series (30, 50, 70, 80, 90, 100%), followed by a replacement step with isoamyl acetate. The samples were then processed through critical point drying (Hitachi-HCP) and coating with gold (Hitachi-MC1000), finally observed with a Hitachi-3400N microscope at the Ultrastructural Microscopy Platform of the IOCAS.

### Collection of nematode materials for sequencing

For all the sequencing in this project, mix-staged *L. marina* of the inbred line F23 were used. Nematodes were cultured on SW-NGM agar plates with OP50 at room temperature (20-25 °C) for a week and harvested using M9 buffer. Samples were washed three times with M9 buffer, following with a 4 h-starvation treatment in M9 buffer on a nutator. After another three times washing with M9 buffer, pelleted samples were then frozen immediately in liquid nitrogen. Genomic DNA was extracted using the CTAB method (Murray and Thompson 1980), total RNA was isolated using the Trizol reagent (Invitrogen, USA), following the manufacturer’s instructions, respectively.

### Genome sequencing

For Illumina short reads sequencing, one library with 350 bp insert size was prepared for paired-end sequencing according to the standard Illumina protocol. Briefly, ∼5 μg genomic DNA (gDNA) was fragmented and size-selected (350 bp) through agarose gel electrophoresis. Selected DNA fragments were blunted and ligated to sequencing adapters. The library was then sequenced on the Illumina NovaSeq platform with a PE150 strategy. Finally, a total of 12.05 Gb (∼65.03×) clean reads were generated after preprocessing. These data were used for assembly polishing and evaluation.

For Pacific Biosciences (PacBio) library construction and sequencing, ∼7 μg gDNA was used to prepare PacBio ∼20 Kb insert size library following the standard PacBio protocol. Two SMRTbell (Single-Molecule Real Time) cells were sequenced on a PacBio Sequel platform (Biomarker Technologies Corporation) with P6-C4 chemistry. 26.11 Gb PacBio data (140.91×) were obtained after preprocessing (including removal of adaptor, low-quality, and reads shorter than 500 bp). 1,989,088 subreads were yielded with an average length of 13.13 Kb and maximum length of 89.55 Kb.

Besides, RNA-Seq sequencing was performed and used for genome annotation. The library was prepared using NEBNext® UltraTM RNA library preparation kit (NEB #E7530) following the manufacturer’s instructions and sequenced on the Illumina platform (HiSeq X Ten) with a PE150 strategy. Raw reads were trimmed for adaptors and low quality. Overall, 9.15 Gb high-quality reads were obtained.

### Genome assembly and evaluation

PacBio subreads were corrected, trimmed and assembled using Canu-based pipeline (http://canu.readthedocs.io/). Briefly, sequencing errors of subreads were corrected using Falcon_sense module embedded in Canu (Koren et al. 2017) (option correctedErrorRate = 0.045). The corrected subreads were trimmed of unsupported bases and hairpin adaptors and then the high-quality PacBio subreads were used for contig assembly. The following parameters were modified: Error correct coverage = 60, GenomeSize = 180 Mb, Asseble coverage = 50 and others leaving as defaults. Totally, 75 contigs were assembled. Finally, base correction was performed with 12.05 Gb Illumina paired-end reads using Pilon (Walker et al. 2014) (v1.22, with parameters --mindepth 10 --changes --threads 4 --fix bases).

To evaluate the quality of the genome assembly, the Illumina short paired-end reads were mapped back to the current assembly applying BWA (Li and Durbin 2009) software (v0.7.10-r789, mapping method: MEM). To assess the completeness of the genome assembly, we used conserved core eukaryotic genes by running software CEGMA (Parra et al. 2007) (v2.5) on the current assembly with the default parameters. We also used the Nematoda conserved genes by running software BUSCO (Simão et al. 2015) (v2.0.1) on the current assembly with the default parameters.

### Gene prediction and functional annotation

Protein coding genes were predicted using a pipeline integrating homology-based prediction, RNA-Seq-based approach, as well as *de novo* prediction (Supplementary Table S7). Briefly, For homology-based gene prediction, protein sequences from *C. elegans* (RefSeq assembly accession: GCF_000002985.6), *C. nigoni* (GenBank assembly accession: GCA_002742825.1), *P. pacificus* (GenBank assembly accession: GCA_000180635.3), and *Brugia malayi* (RefSeq assembly accession: GCF_000002995.3), downloaded from NCBI, were aligned to *L. marina* genome with GeMoMa (Keilwagen et al. 2016) (v1.3.1) with default parameters. For RNA-Seq-based prediction, clean RNA-Seq reads were mapped to *L. marina* genome using Hisat (Kim et al. 2015) (v2.0.4) and assembled by Stringtie (Pertea et al. 2015) (v1.2.3). TransDecoder (Grabherr et al. 2011) (v2.0) and GeneMarkS-T (Tang et al. 2015) (v5.1) were then subsequently applied for the coding region prediction. Transcripts were assembled by Trinity v2.1.1 and then were inputted to PASA (Haas et al. 2003) (v2.0.2) to predict unigenes. For *de novo* annotation, five gene prediction software with default parameters were used, including Genscan (http://hollywood.mit.edu/GENSCAN.html), Augustus (http://augustus.gobics.de/submission), GeneID (Blanco et al. 2007) (v1.4), SNAP (Korf 2004) (v2006-07-28) and GlimmerHMM (Majoros et al. 2004) (v3.0.4). Training models used in Augustus, GlimmerHMM and SNAP were obtained from the above prediction results of PASA (v2.0.2). Finally, gene models from these different approaches were combined by EVidenceModeler (Haas et al. 2008) (EVM, v1.1.1), and later using PASA (v2.0.2) to update EVM annotations to generate UTRs, alternative splicing variation information. After filtering out coding DNA sequences (CDS) shorter than 300 bp, non-triple and with premature stop codon, 17,661 genes were predicted in the genome, which represents v1.0 gene set for *L. marina*.

Functional annotations for predicted genes were performed by BLASTP embedded in BLAST+ package (v2.3.0), with an E value < 1e-5 against a series of nucleotide and protein sequence databases, including NR, KOG, KEGG and TrEMBL. Gene ontology (GO) terms for each gene were assigned by Blast2GO (Conesa et al. 2005) pipeline (v2.5) based on NCBI databases. Approximately 85.39% of all the predicted genes were annotated (Supplementary Table S11).

### Transposable elements analysis

We applied multiple RepeatModeler (v1.0.11) and RepeatMasker (v4.0.9) to identify and classify repetitive sequences in genomes of *L. marina* and *C. elegans*, followed by classification that Wicker *et al*. proposed (Wicker et al. 2007). To determine the details of TEs, we further determined TEs in *L. marina* and *C. elegans* based on methods reported previously by Szitenberg et al. (2016). Then, we conducted comparative study of TEs using results of *L. marina, C. elegans* and data (*C. brenneri, C. remanei, P. pacificus, Ascaris suum, B. malayi, Trichinella spiralis*) from Szitenberg et al. (2016) (https://github.com/HullUni-bioinformatics/Nematoda-TE-Evolution). To obtain more comprehensive insights of long terminal repeat (LTR) retrotransposons, we used combination of LTRharvest and LTRdigest. We also tried homolog-searching method and other methods reported by Kanzaki et al. (2018).

### Gene family identification

Proteins of nine sequenced nematodes were downloaded from Wormbase (version WS270), including *C. elegans, C. nigoni, C. briggsae, C. brenneri, C. remanei, P. pacificus, A. suum, B. malayi, T. spiralis*. OrthoMCL (Li et al. 2003) (v2.0.9; MCL inflation factor 1.5, percent Match Cutoff = 50; e value Exponent Cutoff = −5) was used to identify the orthology. All-against-all BLASTP (Blast+ version 2.3.0) were used to calculate pairwise sequence similarities with P value cutoff of 1e-5 and minimum match length of 50%. In this step, 27,134 gene families for 10 nematodes were identified for further analysis. Number of orthologue groups with all species present, single-copy orthologue groups and species-specific orthologue groups were 1,565, 697 and 7,440, respectively.

### Phylogenetic analysis

In order to determine the phylogenetic position of *L. marina*, 609 core Nematoda lineage orthologues of *L. marina* and nine other nematodes were identified based on BUSCO (version 2.0.1) software. Multiple alignments were performed at the protein level for each of the 609 orthologues using MUSCLE (version 3.8.31, http://www.drive5.com/muscle) and concatenated into one super-gene sequence. RAxML (version 8.2.9) was used to constructed the phylogenetic tree with parameter -f a -m PROTCATJTT and 100 bootstrap replicates were performed for the node support.

### Divergence estimate

MCMCTREE implemented in the PAML (Yang 2007) package (v4.7b) was used to estimate the divergence time. Four calibration time points (*T. spiralis* vs *C. elegans*: 428∼451 MYA, *A. suum* vs *C*.*elegans*: 250∼541 MYA, *P. pacificus* vs *Caenorhabditis* spp: 280∼430 MYA, *C. elegans* vs *C. briggsae*: 5∼30 MYA) were applied based on the TimeTree (http://www.timetree.org) and references (Dieterich et al. 2008; Gordon et al. 2015).

### *Lma-rol-6* gene knockout experiment

sgRNA target sequence of *Lma-rol-6* (ATTCGTCGTCAGAACTACGG) was predicted using CHOPCHOP (https://chopchop.cbu.uib.no/). sgRNA was synthesized via in vitro transcription applying plasmid pDD162 (Addgene, Cat # 47549) and HiScribe™ T7 Quick High Yield RNA Synthesis Kit (New England Biolabs, E2050S), then purified using MEGAclear™ Transcription Clean-Up Kit (Invitrogen, AM1908) as described in the manuals. CAS9 protein was purchased from Clontech (Cat # 632641). For injection, sgRNA and CAS9 ribonucleoprotein complexes were prepared mainly referred to Farboud et al. (Farboud et al. 2019). Specifically, 20 μl injection mixture consisting of 8.75 μM Na-HEPES, pH 7.5, 115 μM KCl, 125 ng/μl CAS9 protein, 25 ng/μl sgRNA, was incubated at 37 °C for 15 min before loading into needles for microinjection. After being injected into gonads, 53 young adult females (P0) were individually transferred to SW-NGM plates and wild type adult males were added for mating. Finally, one roller line was successfully maintained and the mutation was further identified by sequencing.

### Expansion and contraction of gene families

CAFE (De Bie et al. 2006) (version 2.0) was used to infer gene family sizes of ancestral node and analyze gene family expansion and contraction under maximum likelihood framework. The program uses a birth and death process to model gene gain and loss over a phylogeny. The birth and death parameter (λ) was 0.002 and the P value was 0.01.

### Gene synteny analysis

Briefly, predicted protein sequences of *L. marina* were first aligned to *C. elegans* and *C. nigoni* genomes using BLASTP (E value < 1e-5), respectively. Synteny was then detected by MCScanX software (http://chibba.pgml.uga.edu/mcscan2/) with default parameters. A syntenic gene block was defined when at least five homologous gene pairs present, allowing a maximum number of 25 intervening genes/gaps.

### EcR and RXR analysis

We applied the homolog-based method to evaluate the sequence conservation for putative EcRs and RXRs. First, we conducted BLASTP against protein sequences of *Anisakis simplex* (https://parasite.wormbase.org/Anisakis_simplex_prjeb496/Info/Index), *L. marina* and *C. inopinata* (https://parasite.wormbase.org/Caenorhabditis_sp34_prjdb5687/Info/Index), using EcR sequences from *A. summ* (ADY42041.1, ADY42534.1), *B. malayi* (XP_001897313.1, XP_001900768.1), *Haemonchus contortus* (ADD49663.1), *P. pacificus* (ACY82385.1), *Toxocara canis* (KHN78537.1) and *T. spiralis* (XP_003376657.1) as queries. Then, we used hmmscan (v3.1b2, https://www.ebi.ac.uk/Tools/hmmer/search/hmmscan) to search for the DNA-binding domain (DBD, cd07161: NR_DBD_EcR) and ligand binding domain (LBD, cd06938: NR_LBD_EcR) for the putative EcRs of the three nematodes. Taken together, we identified 1 putative EcR in *L. marina* (EVM0014194.1) and *A. simplex* (ASIM_0001371101-mRNA-1), respectively. Although some NHRs in *C. inopinata* showing similarity to EcRs, the LBDs and DBDs of these NHRs are not highly conserved. Further, the identified sequences were aligned with EcRs of *A. suum, B. malayi, Bombyx mori, Drosophila melanogaster, Limulus Polyphemus, P. pacificus, Penaeus vannamei* and *T. spiralis* using MAFFT (v7.205) (Katoh et al. 2002). Phylogenetic tree was build afterwards using FastTree (v2.1.8) (Price et al. 2010) (Amino acid distances: BLOSUM45 Joins: balanced Support: SH-like 1000.Search: Normal +NNI +SPR (2 rounds range 10) +ML-NNI opt-each=1.TopHits: 1.00*sqrtN close=default refresh=0.80. ML Model: Jones-Taylor-Thorton, CAT approximation with 20 rate categories). 1 Putative RXR in *L. marina* (EVM0012017.1) was identified based on BLASTP using RXRs of *Drosophila* sp. (PRF:1614351A), *Dirofilaria immitis* (AAM08268.1), *T. canis* (KHN86367.1), *Wuchereria bancrofti* (EJW88755.1) and *Loa loa* (EFO13605.1) as queries. Phylogenetic trees were constructed using MAFFT and FastTree.

### Positive selection analysis

To identify positive selection genes (PSGs), the branch-site model was used to detect positive selection along the foreground branch. CodeML in PAML (v4.7b) plus a series of likelihood ratio tests (LRTs) were performed on the single-copy orthologs to compute the ratio of synonymous and non-synonymous changes at each codon and test significant differences between the alterative and null models. We conducted a positive selection analysis using the genomic sequences of *L. marina* and nine terristerial nematodes relatives (*C. elegans, C. briggsae, C, nigoni, C, brenneri, C, remanei, P. pacificus, A. suum, B. malayi, T. spiralis*). Branch-site model of the PAML 4 package was used to identify genes with signs of positive selection. As a result, 92 genes possibly under positive selection were identified in the *L. marina* genome (ω > 1, p < 0.05). Of these genes, 49 showed highly significant (p < 0.01) positive selection (Supplementary Table S12).

## DATA ACCESS

The genomic sequence and related datasets of *L. marina* generated in this study have been submitted to the NCBI BioProject database (https://www.ncbi.nlm.nih.gov/bioproject/) under accession number PRJNA577530.

## ACKNOWLEDGEMENTS

We thank members of the L.Z. laboratory for helpful discussions. Thanks also go to Profs. Jianhai Xiang, Fuhua Li, Baozhong Liu, Xiaojun Zhang, Jianbo Yuan’s helps for suggestions, reagents and facilities. The work was supported by the “Yong Scientist Research Program” of Qingdao National Laboratory for Marine Science and Technology (to L.Z.); “Marine life breakthrough funding” of KEMBL, Chinese Academy of Sciences (to L.Z.); National Key R&D Program of China [No. 2018YFD0901301 to Y.X.]; the Marine S&T Fund of Shandong Province for Pilot National Laboratory for Marine Science and Technology (Qingdao) [No. 2018SDKJ0302-1 to L.Z.]; “Talents from overseas Program, IOCAS” of the Chinese Academy of Sciences (to L.Z.); “Qingdao Innovation Leadership Program” [Grant 16-8-3-19-zhc to L.Z.]; and Key deployment project of Centre for Ocean Mega-Research of Science, Chinese Academy of Sciences (to L.Z.). Some nematode strains were provided by the Caenorhabditis Genetics Center, which is funded by NIH Office of Research Infrastructure Programs (P40 OD010440).

## Author contributions

L.Z. and Y.X. designed this project and analysed data. Y.X., P.Z., X.C., Y.S. and H.Y. conducted experiments. L.W., B.X. and X.R. performed the bioinformatics analysis. Y.X., L.Z. and B.X. wrote the manuscript. L.W., P.Z., X.C. and X.R. contributed to manuscript writing and figure preparation. L.Z. supervised the project. All authors read and approved the final manuscript.

## DISCLOSURE DECLARATION

The authors declare that they have no competing interests.

